# A recombinant expression system for the *Plasmodium falciparum* proteasome enables structural analysis of its assembly and the design of selective inhibitors

**DOI:** 10.1101/2025.08.13.670186

**Authors:** Pavla Fajtova, Hanxiao Zhang, Liam Urich, Elany Barbosa da Silva, Cesar Hoffmann da Silva, Jehad Almaliti, Momen Al-Hindy, Evzen Boura, Laura A. Kirkman, Gang Lin, Matthew Bogyo, David A. Fidock, William H. Gerwick, Jianhua Zhao, Anthony J. O’Donoghue

## Abstract

The *Plasmodium falciparum* 20S proteasome (Pf20S) has emerged as a promising antimalarial target. Development of therapeutics to this target has previously relied on native purifications of Pf20S, which is challenging and has limited the scope of previous efforts. Here, we report an effective recombinant Pf20S platform to facilitate drug discovery. Proteasome assembly was carried out in insect cells by co-expressing all fourteen subunits along with the essential chaperone homolog, Ump1. Unexpectedly, the isolated proteins consisted of both a mature and an immature complex. Cryo-EM analysis of the immature complexes revealed structural insights detailing how Ump1 and the propeptides of the β2 and β5 subunits coordinate β-ring assembly, which differ from human and yeast homologs. Biochemical validation confirmed that β1, β2, and β5 subunits of the mature proteasome were catalytically active. Clinical proteasome inhibitors, bortezomib, carfilzomib and marizomib were potent but lacked Pf20S selectivity. However, the tripeptide-epoxyketone J-80 inhibited Pf20S β5 with an IC_50_ of 22.4 nM and 90-fold selectivity over human β5. Structural studies using cryo-EM elucidated the basis for the selective binding of J-80. Further evaluation of novel Pf20S-selective inhibitors such as the reversible TDI-8304 and irreversible analogs, 8304-vinyl sulfone and 8304-epoxyketone, confirmed their potency and selectivity over the human constitutive proteasome. This recombinant Pf20S platform facilitates detailed biochemical and structural studies, accelerating the development of selective antimalarial therapeutics.

## Introduction

The ubiquitin-proteasome system is vital for protein degradation and maintaining cellular balance in eukaryotes, making it an appealing target for new therapies ^1,2^. In human medicine, proteasome inhibitors have achieved remarkable clinical success, particularly in treating multiple myeloma. Drugs like bortezomib, carfilzomib, and ixazomib have transformed patient care ^3-6^. This success has spurred research into using proteasome inhibition as a strategy to combat parasitic infections. Recent studies have identified significant structural and functional differences between human and parasitic proteasomes, creating opportunities for developing selective inhibitors ^7-10^. Notable progress has also been made in targeting the proteasomes of protozoan parasites such as *Leishmania donovani* and *Trypanosoma cruzi*. Pharmaceutical companies such as GSK and Novartis are developing parasite-specific proteasome inhibitors with minimal cross-reactivity to the human enzyme ^11,12^.

*Plasmodium falciparum* (Pf), the causative agent of the most lethal form of malaria, relies heavily on its proteasome for development and survival throughout its complex life cycle ^13,14^. The *P. falciparum* 20S proteasome (Pf20S) has been recognized as a promising drug target, with pioneering work by multiple research groups establishing its potential.

It has been demonstrated that parasite proteasome activity is critical during the intraerythrocytic stages, and selective inhibitors with antimalarial activity have already been identified ^7,15-17^. In addition, several natural products and their derivatives have been discovered that inactivate Pf20S ^18,19^. Importantly, proteasome inhibitors can overcome resistance to frontline antimalarials ^20^. One particularly attractive feature is that proteasome inhibitors synergize with artemisinin derivatives, which are the cornerstone of all first-line antimalarial combination therapies ^7^. This synergy is also observed in *P. falciparum* parasites that have acquired partial resistance to artemisinins, a substantial clinical problem in Asia that has now also emerged in multiple African countries ^21-23^. The mechanistic basis of this interaction is thought to arise from the critical role of the 20S proteasome in eliminating parasite proteins damaged by the alkylating action of activated artemisinins ^24^. Inhibiting the proteasome’s capacity to clear artemisinin-damaged proteins is therefore a highly appealing strategy to sustain artemisinin efficacy and to develop potent combination therapies that incorporate a *P. falciparum*-specific proteasome inhibitor.

While most inhibitors directly interact with the substrate binding pocket and catalytic threonine, others have been discovered to bind at adjacent sites ^7,25^. Collectively, these studies underscore the importance of Pf20S as a therapeutic target against malaria. Yet despite the promising nature of Pf20S as a drug target, biochemical and structural studies have been hindered by the difficulty in obtaining sufficient quantities of the highly purified native enzyme. Recombinant expression systems using insect cells have successfully overcome similar challenges for human, *Trichomonas vaginalis*, and *Trypanosoma cruzi* proteasomes ^26-28^. These systems have produced fully functional proteasomes, facilitating structural and functional studies that advance drug discovery. However, the recombinant expression of a functional Pf20S has remained elusive.

In this study, we successfully generated a recombinant Pf20S proteasome by co-expressing all fourteen constituent subunits (seven α and seven β) along with the *P. falciparum* Ump1 chaperone in an insect cell expression system. The Ump1 chaperone proved crucial for proper assembly, as no functional enzyme complex was detected without co-expression of the chaperone, consistent with findings in other organisms ^27,29,30^. The recombinant proteasome exhibits biochemical and structural properties identical to those of the native enzyme isolated from parasite cultures. This recombinant enzyme enabled the discovery of a β5-selective substrate and allowed for extensive inhibition studies with a diverse panel of compounds. Furthermore, structural studies revealed how a tripeptide-epoxyketone inhibitor binds with high affinity to the β5 subunit. Overall, the recombinant expression system overcomes the limitations associated with native proteasome isolation and establishes a platform for detailed investigation into the structure and function of protozoan proteasomes ^31,32^. This advancement will accelerate the development of novel antimalarial therapeutics with reduced host toxicity.

## Results

### Assembly and isolation of recombinant Pf20S

The *P. falciparum* proteasome is a validated target for antiparasitic drug development. However, a key bottleneck exists in isolating Pf20S from parasite extracts. The published purification protocols involve two to five time-consuming chromatography steps, yielding limited protein quantities that are only sufficient for small scale biochemical or structural studies ^7,9,17,25^ This necessitates regular isolation of fresh proteasome, which in turn requires continuous culturing of the parasite. For other anti-microbial drug targets, the availability of recombinant proteases has facilitated high-throughput screening, concentration-response studies or structural studies ^33-35^. Therefore, we set out to establish a recombinant platform for Pf20S. Encouragingly, our recent work demonstrated that the *Trichomonas vaginalis* proteasome (Tv20S) can be successfully expressed in an Sf9 insect cell system ^27^, paving the way to consider additional parasite proteasome targets.

The genome of *P. falciparum* encodes seven α and seven β subunits of Pf20S, all of which align with homologous subunits in the human constitutive proteasome (**Fig. S1**). The challenge in generating recombinant Pf20S proteasomes lies in the assembly of the different subunits, which need to be precisely arranged in the correct order. Initial attempts of expression and purification showed that only the proteasome from the host insect cells, *Spodoptera frugiperda* 20S proteasome (Sf20S) was detectable by native PAGE when imaged using the fluorogenic activity-based probe, Me4BodipyFL-Ahx3Leu3VS (**Fig. 1a**). These studies indicated that assembly and maturation of recombinant Pf20S was unable to occur using the host chaperone proteins. We and others have previously highlighted the importance of the chaperone protein Ump1 for successful expression of a fully functional proteasome ^27,29,30^. Using protein alignment searches, we identified a homolog of Ump1 in the *P. falciparum* genome (Uniprot ID: A0A5K1K941). Co-expression of this gene with the α and β subunits successfully produced the recombinant 20S proteasome as detected by native gel electrophoresis using the activity-based probe. Furthermore, it could be clearly differentiated from the host proteasome by its different migration pattern on the native gel.

**Figure 1.**
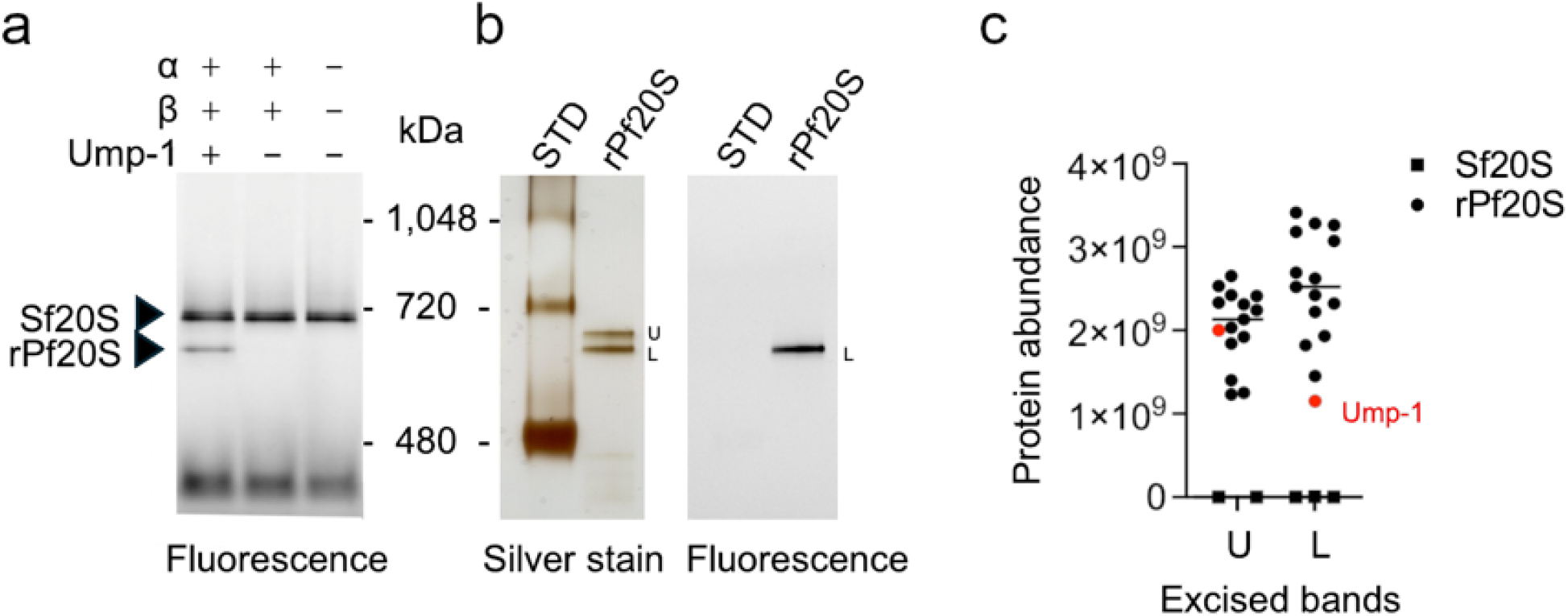
Characterization of recombinant Pf20S proteasome. **a**. Native-PAGE analysis of insect cell lysates following baculovirus co-infection, labeled with Me4BodipyFL-Ahx3Leu3VS fluorescent activity-based probe. **b**. Native-PAGE gel of purified recombinant Pf20S proteasome. Gels were visualized by silver staining (total protein) and fluorescent scanning (active proteasome complexes). STD is a protein standard. **c**. Proteasome subunit abundance in the upper (U) and lower (L) excised gel bands. Detailed proteomic data are provided in the Source Data file.

Recombinant Pf20S contains a C-terminal twin-strep tag on the β7 subunit and was therefore selectively enriched from the cell lysates using a streptavidin column. Unexpectedly, analysis of the silver-stained native gel after streptavidin enrichment revealed two protein complexes that had a similar migration pattern (**Fig. 1b**). Both complexes were distinct from the insect proteasome. Upon addition of the activity-based probe only the lower band was labelled. These findings suggest that the lower band represents the mature, functional Pf20S proteasome, whereas the upper band likely corresponds to an immature proteasome complex. We attempted to separate these complexes by size exclusion chromatography, but they eluted as a single peak.

We hypothesized that the difference between the upper and lower band could be attributed to the retention of propeptides from the β1, β2, and β5 subunits and the retention of the Ump1 chaperone which would change the overall molecular weight and charge of the complex. To test this, the upper and lower bands were excised from the silver-stained gel (**Fig. 1b**) and subjected to proteomics analysis by searching against the *S. frugiperda* and *P. falciparum* proteome. In each band, the seven α and seven β subunits of Pf20S were amongst the most abundant proteins present (**Fig. 1c**). However, Ump1 was lower in abundance in the lower band indicating that it may have been degraded and released from the matured 20S proteasome. Several insect proteasome subunits were also found but their peak intensities were more than 1000-fold lower in abundance than the subunits of *P. falciparum* (**File S1**), therefore ruling out the presence of appreciable levels of chimeric insect-*Plasmodium* proteasomes. We next looked at the peptides derived from the propeptide region of the catalytic subunits to determine if they were higher in abundance in the upper band relative to the lower band. For all three catalytic subunits, the propeptide sequences were 1.8-fold to 4.8-fold more abundant in the upper band while the average intensity for peptides derived from the mature region of the subunits showed no difference (**File S1**). These studies strongly indicate that the upper band consists of an immature proteasome complex that retains the propeptide sequences and higher amounts of Ump1. When considering both the upper and lower bands together, a yield of 1 mg per liter at >98% purity was achieved for recombinant Pf20S using the insect expression system.

### Structural evaluation of immature 20S proteasomes

To gain structural insights into both the mature and immature proteasomes, we analyzed the recombinant Pf20S using single particle cryogenic electron microscopy (cryo-EM). Two-dimensional classification of the cryo-EM images revealed the presence of both correctly and incorrectly assembled complexes (**Fig. S2**). In human cells, normal proteasome assembly involves the dimerization of two half-proteasomes through the association of their β-rings, yielding the nascent core particle. This intermediate is subsequently remodeled by the stepwise removal of the N-terminal propeptides of β1, β2 and β5 followed by the degradation of Ump1 ^30,36-39^. The incorrectly assembled Pf20S complexes consist of two half-Pf20S sub-complexes that are misaligned at the β-β dimerization interface. Three-dimensional refinement of the misaligned complexes resulted in a cryo-EM map where one half-Pf20S displayed high-quality density at ∼3.0 Å resolution, while the other half-Pf20S showed low-quality cryo-EM density (**Fig. 2a**). This suggests that the two half-Pf20S sub-complexes are misaligned in multiple configurations, resulting in blurring of the cryo-EM density after averaging. Analysis of the high-resolution half-Pf20S revealed clear density for Ump1, in addition to the propeptides of β2 and β5 within the antechamber (**Fig. 2b**). However, no density for the β1 propeptide was observed in the cryo-EM map.

**Figure 2.**
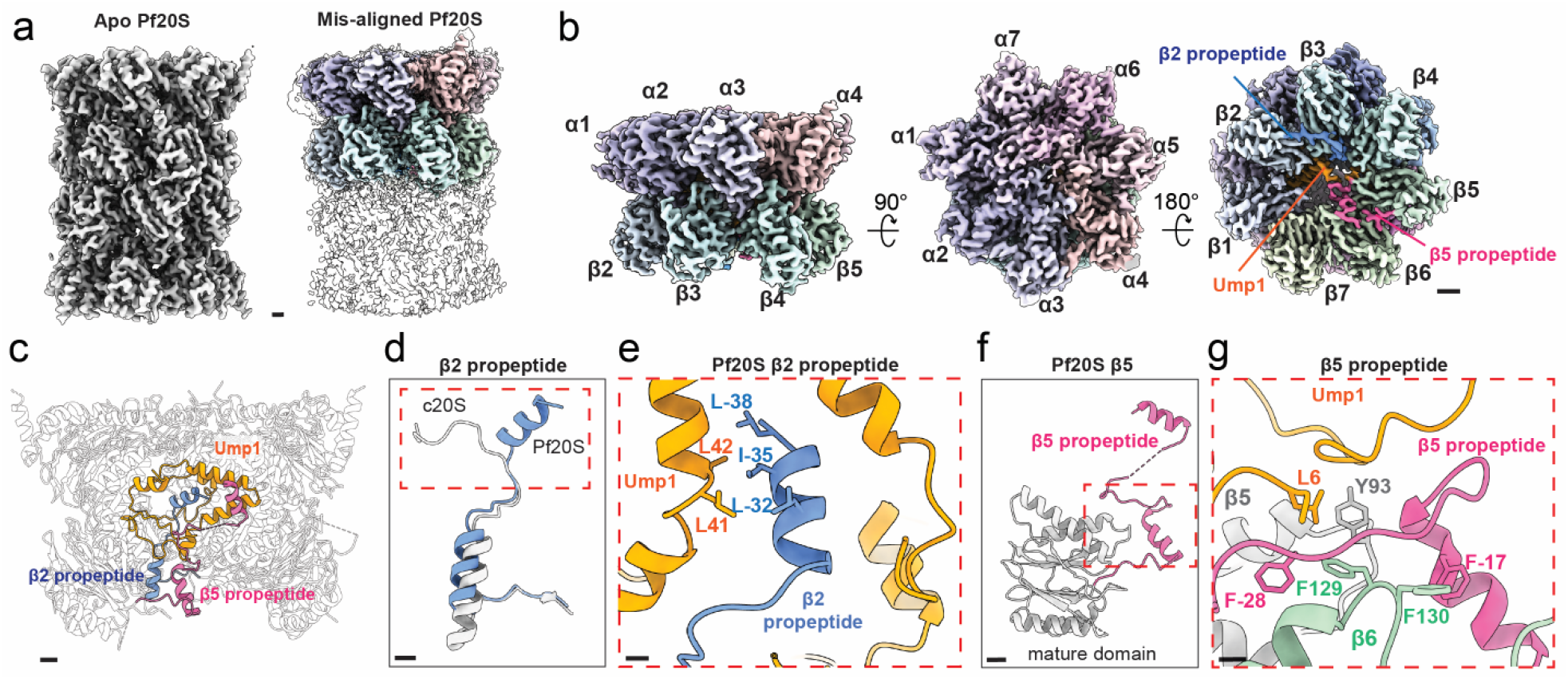
Incorrectly assembled half-Pf20S displays distinct β2 and β5 propeptides compared to the human 20S. **a**. Cryo-EM map of incorrectly assembled half-Pf20S, showing top-down view and bottom-up view. Scale bar, 13 Å. **b**. Atomic model of the half Pf20S, illustrating Ump1 and β2 and β5 propeptides positioned within the antechamber formed by the α-ring and β-ring. Scale bar, 13 Å. **c**. Interaction between Pf20S Ump1 and N-terminal helix of β2 propeptide. Scale bar, 5 Å. **d**. Overlay of β2 propeptides from human 20S (grey) (PDB ID: 7NAN) and Pf20S (blue) (PDB ID: 9Y0L). Red box highlights the N-termini of the two β2 propeptides, which adopt different conformations. Scale bar, 10 Å. **e**. Stabilization of the β2 propeptide by Ump1. Scale bar, 20 Å. **f**. Atomic model of Pf20S β5 (grey) wth the propeptide region highlighted (pink). Red box highlights the N-terminal region of β5 propeptide. Scale bar, 10 Å. **g**. An α helix (pink) within the N-terminal region of the β5 propeptide interacts with the β6 (green) and β5 (grey) subunits. Scale bar, 20 Å.

The structure of the half-Pf20S reveals how the β2 and β5 propeptides cooperate with Ump1 to coordinate β-ring assembly. Ump1 adopts a conformation similar to the human homolog POMP ^40,41^. Meanwhile, the *Plasmodium* β2 and β5 propeptides show structural differences compared to their mammalian and yeast counterparts ^42^. In the human proteasome, the C-terminus of the β2 propeptide forms a loop that points toward the wall of the antechamber ^40,43^, while the C-terminus of the Pf20S β2 propeptide adopts an α-helical conformation and points toward the center of the antechamber (**Fig. 2c-d**). This α-helix is stabilized by hydrophobic interactions involving Leu41 and Leu42 of Ump1 and Leu(-32), Ile(-35), and Leu(-38) of the β2 propeptide (numbering corresponding to the mature sequence) (**Fig. 2e**). Our structure also provides a nearly complete model of the β5 propeptide, shedding light on how it facilitates β-ring assembly. Starting from the catalytic threonine (Thr1), the β5 propeptide adopts a loop-helix-loop-helix conformation that is similar to the β2 propeptide (**Fig. 2f**). The N-terminal α-helix is stabilized by π-stacking between Phe(-17) of β5 and Phe130 of β6. Additionally, the loop between the two α-helices in the β5 propeptide is sandwiched between Ump1 and β6 and is stabilized by stacking of β5 Phe(-28), β6 Phe129, and β5 Tyr93 to form a hydrophobic pocket that is occupied by Ump1 Leu6 (**Fig. 2g**). These results reveal how the β5 propeptide coordinates with Ump1 to recruit β6 during β-ring assembly.

Comparing the structure of the incorrectly assembled recombinant Pf20S with the native enzyme (PDB ID: 7LXU) offers insights into the factors contributing to the incorrect assembly. In the native enzyme, the C-terminal tails of the β1 and β7 subunits coordinate the dimerization of two half-proteasomes by extending across the dimerization interface to bind the opposing half-20S complex ^40,43^. However, in the incorrectly assembled recombinant Pf20S the C-terminus of β1 remains unbound (**Fig. S3**). The absence of this important, dimer-stabilizing interaction likely reduces the binding affinity between the two proteasome halves, leading to an unstable complex. The incorrectly assembled Pf20S complexes are enzymatically inactive as the catalytic subunits could not be labelled by the activity-based probe. Due to their inactivity, they do not interfere with downstream biochemical assays. Using the silver-stained gel (**Fig. 1b**), we determined that these inactive complexes (upper band) represent 37% of the total Pf20S protein. This allowed us to accurately calculate the concentration of active enzyme associated with the lower band. To confirm that this active recombinant enzyme is a suitable substitute for the native form, we performed a series of biochemical studies.

### Biochemical validation of recombinant Pf20S

To assess subunit activity, the purified recombinant enzyme was incubated with the activity-based probe, and the labeled subunits were visualized on a denaturing protein gel. Previous studies had established that this probe selectively labels only the β2 and β5 subunits of the native enzyme, therefore, only two bands were anticipated to be seen for the recombinant enzyme ^15^. Indeed, two distinct bands appeared at 25 and 23 kDa, confirming the functionality of the β2 and β5 subunits (**Fig. 3a**). Our previous studies revealed that the tripeptide epoxyketone inhibitor, J-80, is a potent and specific inhibitor of the native Pf20S β5 subunit ^9^. Preincubation of the recombinant enzyme with J-80 prior to labeling active subunits with the probe resulted in the elimination of the lower band, confirming that the lower band was indeed the β5 subunit. In a parallel experiment, the enzyme was preincubated with Ac-Phe-Arg-Ser-Arg-epoxyketone (FRSR-ek), a tetrapeptide inhibitor developed by our group to inhibit the β2 subunit of *T. vaginalis* ^44^. This inhibitor was predicted to bind the Pf20S β2 subunit due to the presence of Arg in the P1 position. Indeed, FRSR-ek decreased the labeling of the upper β2 band with the probe. As a control, the broad-spectrum inhibitor marizomib (MZB) eliminated the labeling of both bands. To confirm β1 subunit activity, recombinant Pf20S was assayed using a previously established fluorogenic substrate: the acetylated tetrapeptide Ac-Nle-Pro-Nle-Asp linked to a C-terminal 7-amino-4-methylcoumarin fluorescent reporter group (Ac-nPnD-amc). Pf20S-mediated cleavage of this substrate was inhibited by the broad-spectrum inhibitor MZB (which targets all subunits), but not by the selective β5 inhibitor J-80 or the selective β2 inhibitor FRSR-ek (**Fig. 3b**) confirming that Ac-nPnD-amc was detecting β1 activity and not β2 or β5. In summary, a combination of the gel-based assays and the fluorescent plate assay confirmed that the β1, β2, and β5 subunits are functionally active in the recombinant Pf20S enzyme.

**Figure 3.**
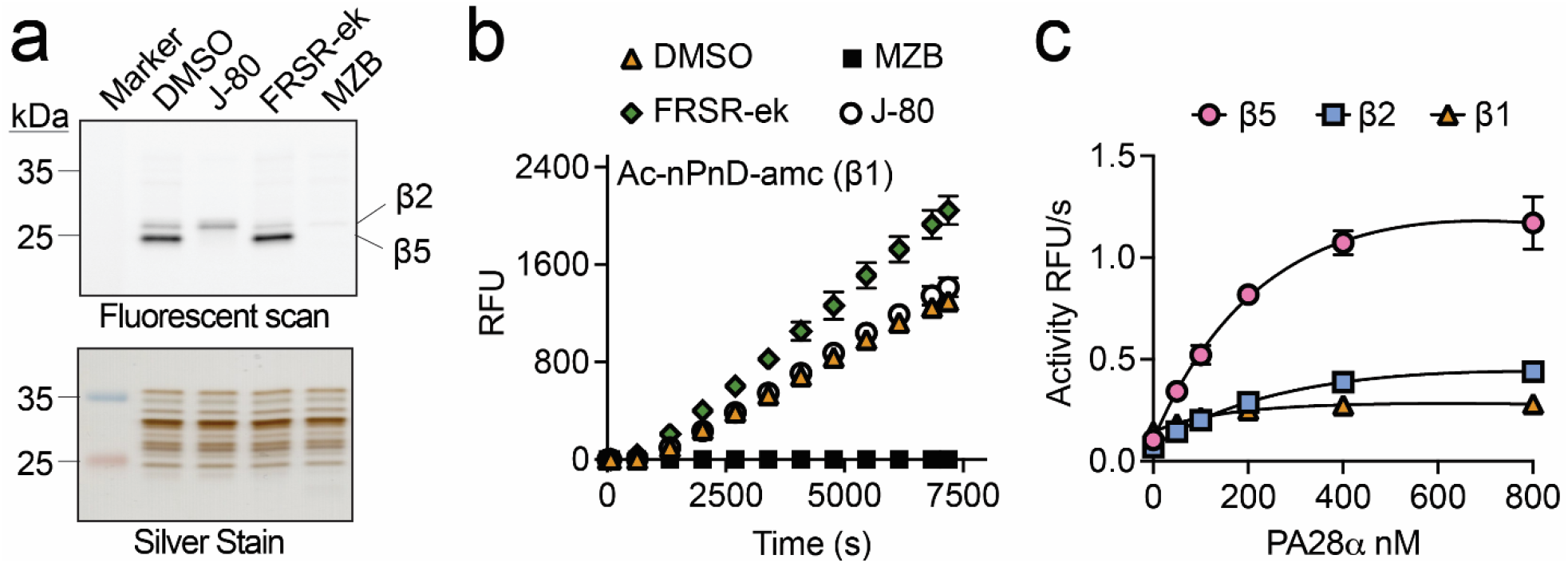
Biochemical characterization of proteasome activity. **a**. SDS-PAGE analysis of proteasome samples. Upper panel: fluorescence scan of protein gel labelled with Me4BodipyFL-Ahx3Leu3VS activity-based probe revealing the β2 and β5 catalytic subunits. Lower panel: silver-stained gel showing molecular weight marker and equal protein loading in each lane. **b**. Time-course analysis of proteasome caspase-like (β1) activity using Ac-nPnD-amc fluorogenic substrate. Activity measured in the presence of proteasome inhibitors. Data shown as the change in relative fluorescence units (RFUs) over time. **c**. Concentration-response curves showing the effect of PA28α regulatory subunit on proteasome activity. Different proteasome subunits (β5, β2, β1) show distinct responses to increasing PA28α concentrations (0-800 nM). Assays were performed in technical replicates (n□=□3).

To ensure a complete set of subunit-specific substrates was available for microwell plate assays with the recombinant Pf20S enzyme, we evaluated the previously published reporter substrates for β2 (Z-VLR-amc) and for β5 (Suc-LLVY-amc and Ac-WLA-amc). Although these substrates were efficiently cleaved, a key difference emerged: activity with Z-VLR-amc (the β2 substrate) was effectively inhibited by FRSR-ek (a β2 inhibitor), whereas Suc-LLVY-amc and Ac-WLA-amc (the putative β5 substrates) were not completely inhibited by the β5 inhibitor, J-80 (**Fig. S4**). This finding indicates that Suc-LLVY-amc may be cleaved by more than one proteasome subunit. Consequently, we evaluated a new β5 substrate, Ac-Phe-Asn-Lys-Leu-amc (Ac-FNKL-amc), which was recently developed for assaying the *Schistosoma mansoni* 20S proteasome ^45^. We found that Pf20S efficiently cleaved Ac-FNKL-amc. Importantly, this substrate proved specific for the β5 subunit, as its cleavage was completely inhibited by J-80. The K_M_ value was calculated for the β1 (Ac-nPnD-amc), β2 (Z-VLR-amc) and β5 (Ac-FNKL-amc) substrates using recombinant Pf20S (**Fig. S4**).

Previous studies have shown that human proteasome activator subunit 1 alpha (PA28α) increases enzyme activity of the native Pf20S ^8^. To assess the effect of PA28α on the activity of each catalytic subunit of recombinant Pf20S, fluorogenic peptide substrates specific for β1 (Ac-nPnD-amc), β2 (Z-VLR-amc), and β5 (Ac-FNKL-amc) were employed. With fixed concentrations of enzyme and substrate, assays were performed in the presence of increasing PA28α concentrations up to 800 nM. Subunit activity increased linearly as PA28α concentration increased from 0 to 200 nM, with a less pronounced increase between 200 and 800 nM (**Fig. 3c**). PA28α is known to bind to the α-rings of the proteasome, which enhances substrate access to the catalytic core. At a concentration of 200 nM PA28α, the activity of the β5 subunit increased 8-fold, while β1 and β2 subunit activities increased 4-fold and 2-fold, respectively. Taken together these studies indicate that the α-ring of the recombinant enzyme is correctly assembled as it binds to PA28α; and a 1 nM to 200 nM ratio of recombinant Pf20S to PA28α is ideal for downstream enzyme activity assays.

### Assessment of clinical proteasome inhibitors as potential anti-malarial inhibitors

Given the clinical success of proteasome inhibitors like carfilzomib and bortezomib in treating multiple myeloma, and the clinical trials of the brain-penetrable inhibitor marizomib for glioblastoma, we aimed to directly compare the potency of these compounds against both Pf20S and human constitutive 20S (c20S) proteasomes. Although these inhibitors are known to be toxic to human cells, our goal was to determine if any could serve as a viable starting point for a dedicated anti-malarial drug development program. Bortezomib primarily targeted the β5 subunits of both c20S (IC_50_ = 5.2 ± 0.5 nM) and Pf20S (IC_50_ = 1.9 ± 0.3 nM), and it also inhibited the β1 and β2 subunits of both enzymes, albeit at higher concentrations (**Fig. 4a, Table 1**). Carfilzomib inhibited both the β5 (IC_50_ = 6.5 ± 0.4 nM) and β2 (IC_50_ = 13.4 ± 0.8 nM) subunits of Pf20S, but it also targeted the c20S β5 subunit (IC_50_ = 15.2 ± 1.9 nM) within a similar concentration range (**Fig. 4b**) Finally, marizomib inhibited the β5 subunits of both Pf20S and c20S with IC_50_ values in the 8-20 nM range, while also targeting both β2 subunits (Pf20S and c20S) and the c20S β1 subunit with IC_50_ values of ∼300 nM (**Fig. 4c**). Collectively, these findings demonstrate that the clinical proteasome inhibitors tested exhibit no significant selectivity for Pf20S over c20S, which therefore makes them unsuitable as direct anti-malarial therapies. Consequently, we suggest that they are not useful starting points for specific anti-malarial drug development efforts.

**Table 1:** IC_50_ values of clinical proteasome inhibitors and Plasmodium-selective inhibitors against Pf20S. Assays were performed in three technical replicates, and the values are reported in nM +/- SD. Compounds that do not inhibit enzyme activity at 8 μM are noted.

| Inhibitor | β5 Pf20S | β2 Pf20S | β1 Pf20S | β5 c20S | β2 c20S | β1 c20S |
| --- | --- | --- | --- | --- | --- | --- |
| **Bortezomib** | 1.97 ± 0.31 | >3000 | 241.80 ± 80.99 | 5.17 ± 0.49 | >3000 | 88.06 ± 8.74 |
| **Carfilzomib** | 6.50 ± 0.42 | 13.43 ± 0.83 | no inhibition | 15.21 ± 1.93 | 195.60 ± 20.37 | 1210 ± 177 |
| **Marizomib** | 8.83 ± 0.81 | 356.50 ± 52.48 | >8000 | 19.28 ± 1.12 | 236.10 ± 19.99 | 391.20 ± 39.52 |
| **J-80** | 22.35 ± 1.27 | >8000 | no inhibition | 832.10 ± 63.69 | no inhibition | no inhibition |
| **TDI-8304** | 3.10 ± 0.35 | no inhibition | no inhibition | no inhibition | no inhibition | no inhibition |
| **EY-4-78** | 11.98 ± 0.72 | no inhibition | no inhibition | 774.70 ± 76.95 | >8000 | no inhibition |
| **8304-vs** | 3.47 ± 0.21 | >8000 | no inhibition | 852.80 ± 117.80 | no inhibition | no inhibition |
| **8304-ek** | 23.67 ± 2.06 | 2159 ± 908 | no inhibition | 261.20 ± 32.42 | >8000 | no inhibition |

**Figure 4.**
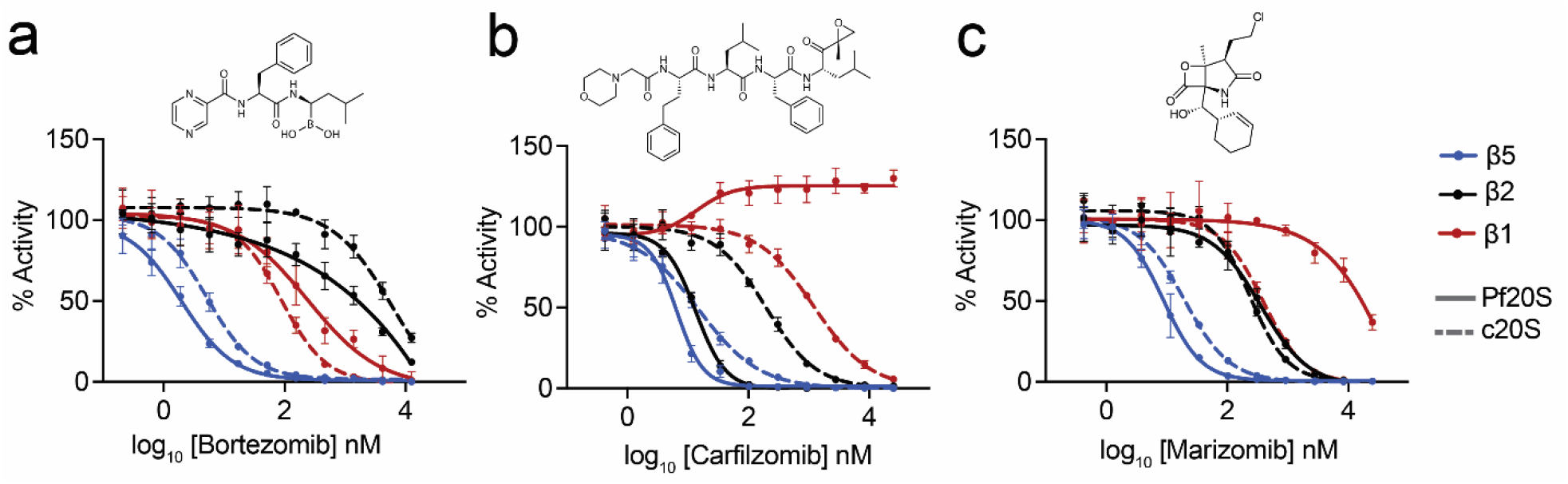
Testing clinical proteasome inhibitors. Chemical structures and concentration response assays for **a**. bortezomib, **b**. carfilzomib and **c**. marizomib against Pf20S (solid lines) and human c20S (dashed lines). Assays were performed in technical triplicate reactions using β5, β2 and β1 specific substrates. Assays were performed in technical replicates (n□=□3).

### J-80 inhibition of Pf20S β5 revealed by structural studies

Previous studies established J-80 as a potent inhibitor of the Pf20S β5 subunit, demonstrating 2,500-fold selectivity in killing *P. falciparum* cultures (IC_50_ = 9.2 ± 1.8 nM) compared to mammalian cell cultures (IC_50_ = 24,300 ± 2,300 nM) ^9^. Using our recombinant Pf20S, we confirmed the specificity of J-80 for the β5 subunit (IC_50_ = 22.4 ± 1.3 nM), observing no inactivation of the β2 or β1 subunits at concentrations up to 35 μM (**Fig. 5a, Table 1**). Direct comparison of J-80 activity against the human constitutive 20S proteasome (c20S) further revealed that it also targeted the c20S β5 subunit (IC_50_ = 832 ± 63.7 nM) but with 90-fold lower potency than Pf20S β5.

**Figure 5.**
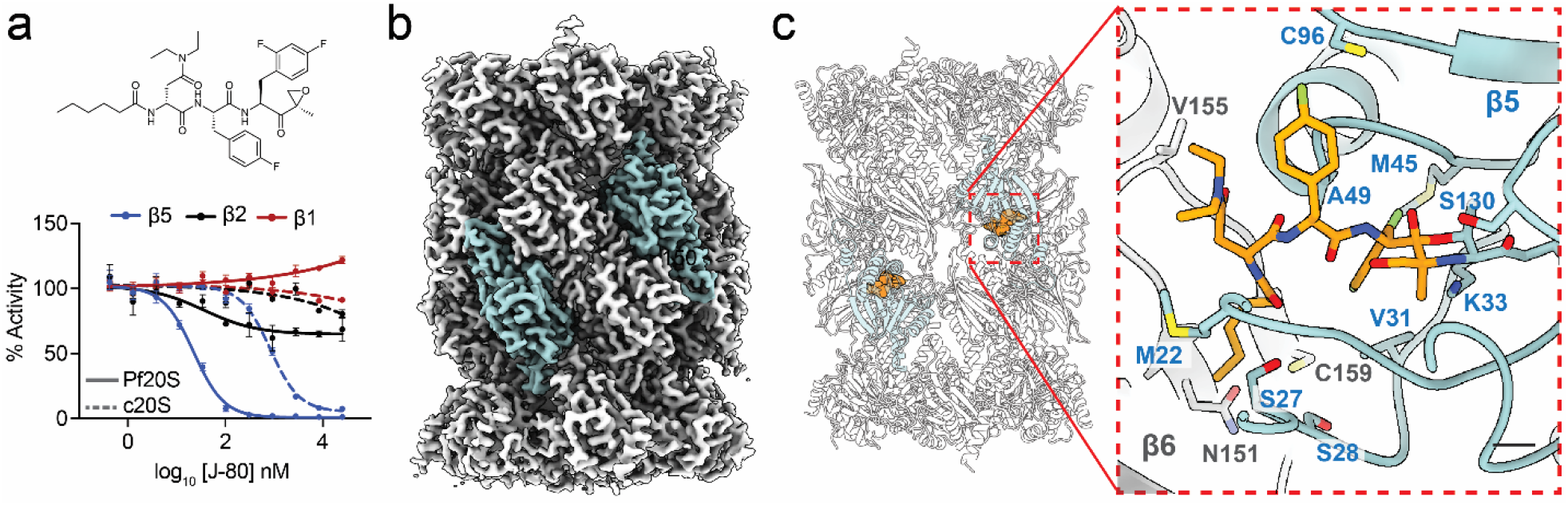
J-80 covalently interacts with the β5 substrate binding pocket of Pf20S. **a**. Structure of J-80 and concentration response assay against Pf20S (solid lines) and human c20S (dashed lines). Assays were performed in technical triplicate reactions using β5, β2 and β1 specific substrates. **b**. The cryo-EM density map with the β5 subunit highlighted in light blue. Scale bar, 10 Å. **c**. Ribbon structure of Pf20S. In the inset, J-80 (orange) (PDB ID: 9Y0K) is bound to the β5 catalytic Thr1, forming a ring. Interactions occur between J-80 and the substrate binding pocket of β5 (light blue ribbon) and β6 (grey ribbon). Scale bar, left: 10 Å, right: 2.5 Å. Enzymatic assays were performed in technical replicates (n□=□3)

To elucidate the structural basis for J-80’s differential potency against Pf20S β5 versus c20S β5, we determined the structure of J-80 bound to Pf20S by cryo-EM. For this analysis, only correctly assembled mature Pf20S complexes were examined, as J-80 cannot bind to immature complexes lacking functional active sites. Analysis of the cryo-EM data yielded a ∼3.0 Å resolution map that shows clear continuous electron density extending from the catalytic threonine (Thr1) of the β5 subunit that resembles J-80. This confirmed that the inhibitor is bound to the β5 subunit (**Fig. 5b**). Consistent with its biochemical specificity, no substantial electron density was detected in the substrate-binding pockets of the β1 and β2 active sites. Comparison between the apo-Pf20S structure (unbound) and the J-80/Pf20S complex showed good structural alignment, with no global conformational changes induced by J-80 binding (**Fig. S5**).

J-80 is coordinated by multiple residues within the substrate-binding groove at the interface of the β5 and β6 subunits. A ring forms between Thr1 of β5 and the epoxyketone warhead of the inhibitor ^27^. This ring product is stabilized by interactions with Lys33 and Ser130 of the β5 subunit (**Fig. 5c**). The 2,4-difluorobenzene ring at the P1 site of J-80 occupies a hydrophobic pocket formed by Val31, Met45, and Ala49 of β5, while the P2 mono-fluorobenzene ring coordinates with Cys96 of β5. The P3 *D*-diethyl-asparagine substituent of J-80 is positioned between Met22 of β5 and Val155 of β6, and it also interacts with structural features of the P2 binding pocket. Furthermore, the P4 hexanoyl group of J-80 tucks into a pocket formed by Ser27 and Ser28 of β5, and a β-sheet region of β6 (spanning Asn151 to Cys159). Additionally, the peptide-backbone of J-80 forms multiple hydrogen bonds along the substrate binding groove, interactions that likely stabilize its binding and contribute to its high potency and specificity for Pf20S β5. This structural data further validates the recombinant Pf20S as a suitable platform for therapeutics development. In addition, it highlights J-80 as an ideal starting point for structure-based inhibitor development.

### Validation of Pf20S-selective inhibitors

Our findings with J-80 demonstrated that targeting the Pf20S β5 subunit was sufficient to kill *P. falciparum*. Consequently, we evaluated additional Pf20S-specific proteasome inhibitors that were developed by colleagues at other institutes. Our initial assessment focused on EY-4-78, a tripeptide-vinyl sulfone inhibitor. This compound was optimized through medicinal chemistry from an earlier generation of Plasmodium-selective tripeptide-vinyl sulfone inhibitors ^7^ to achieve improved selectivity, solubility, and oral bioavailability ^46^. Gel-based competition assays using a fluorescent activity-based probe indicated that EY-4-78 was β5-selective; however, the exact potency of this compound had not yet been determined. In this study, we revealed that EY-4-78 has an IC_50_ of 12.0 ± 0.7 nM for Pf20S β5 and exhibits 65-fold selectivity over c20S β5, although it also targets Pf20S β2 in the micromolar range (**Fig. 6a, Table 1**). We also assessed TDI-8304, the lead compound emerging from another medicinal chemistry program aimed at improving the solubility, permeability, and metabolic stability of an initial hit from a series of macrocyclic proteasome inhibitors, while maintaining potency ^47^. TDI-8304 was identified in this study as a highly potent and specific inhibitor of Pf20S β5 with IC_50_ = 3.1 ± 0.4 nM (**Fig. 6b**) with no detected inhibition of c20S subunits at 12.5 μM, the highest concentration tested. TDI-8304 binds to the β5 subunit with the P1 cyclopentyl group projecting into the S1 pocket and the P3 morpholino group inserting into the S3 pocket ^20^ (**Fig. 6c**). These interactions are distinct from J-80 that binds to the S1 to S4 pockets (**Fig. 6d**). Although these two compounds are structurally distinct, a key differentiating feature is their mode of inhibition. J-80 binds irreversibly to the β5 subunit by covalently interacting with the catalytic Thr1 nucleophile (**Fig. 6e**), while TDI-8304 binds reversibly to the binding pocket without directly engaging the catalytic residue (**Fig. 6f**).

**Figure 6:**
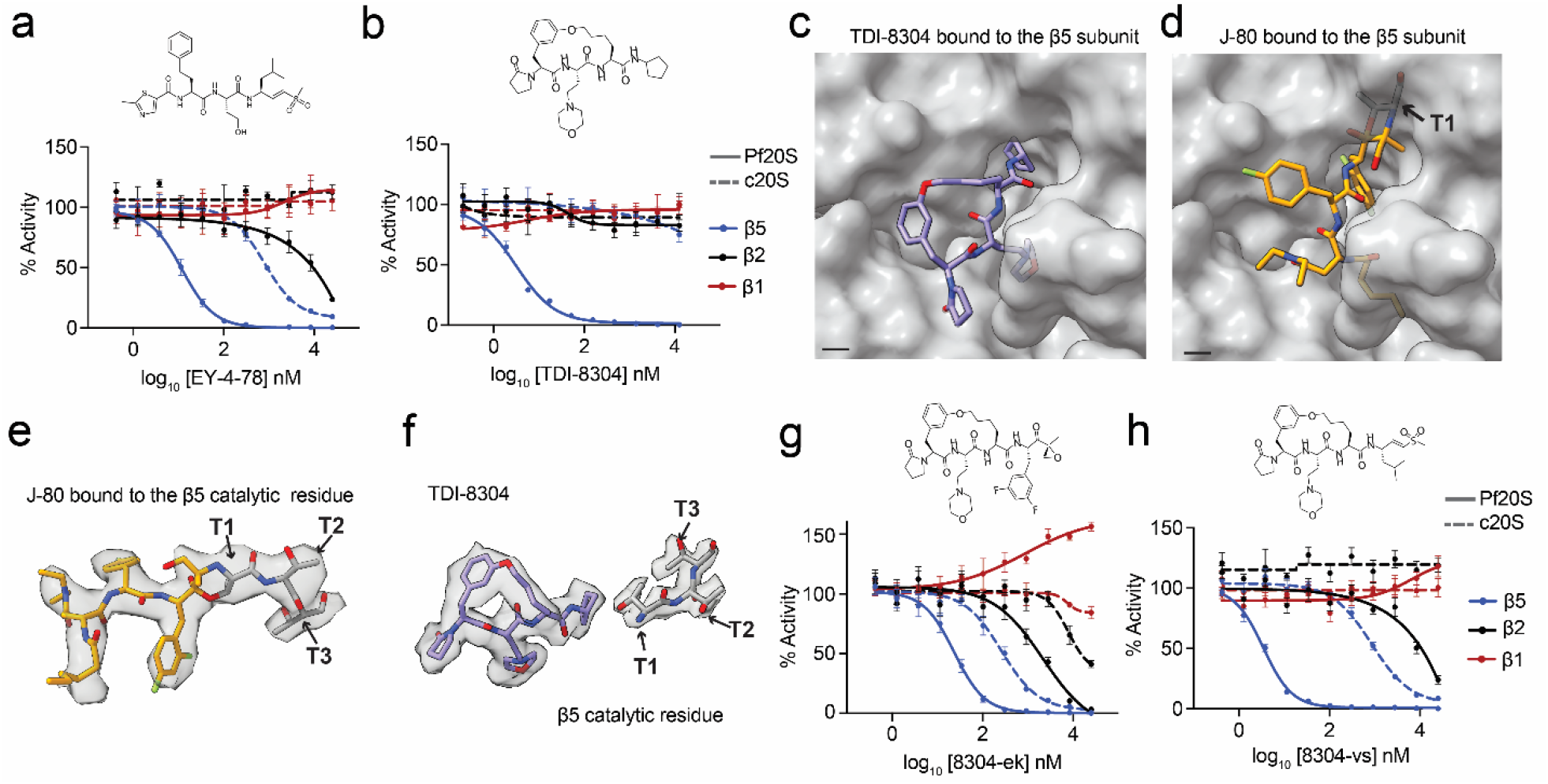
Development of Pf20S-selective inhibitors. Chemical structure and concentration response assays with **a**. EY-4-78 and **b**. TDI-8304. **c**. Structure of TDI-8304 bound to the β5 subunit of Pf20S (PDB ID: 8G6E). Scale bar, 2.5 Å. **d**. Structure of J-80 bound to the β5 subunit of Pf20S (PDB ID: 9Y0K). Scale bar, 2.5 Å. **e**. Irreversible binding of J-80 to Thr1. **f**. TDI-8304 binds reversibly to β5 but does not directly interact with Thr1. Chemical structure and concentration response assays with **g**. 8304-epoxyketone and **h**. 8304-vinyl sulfone. All assays were performed in technical triplicate reactions using β5, β2 and β1 specific substrates.

### Evaluation of irreversible TDI-8304 analogs

Comparative studies using proteasome mutant lines revealed that TDI-8304 exhibited the highest shift in IC_50_ values when compared to J-80, and EY-4-78, also an irreversible binding inhibitor ^46^. It was postulated that for reversible inhibitors, a mutation-induced reduction in target residence time could be a significant disadvantage, as an irreversible inhibitor might eventually overcome weaker binding affinity to a mutant enzyme if given sufficient time.

Based on these observations, we developed two hybrid inhibitors that combined the macrocyclic scaffold of TDI-8304 with a P1-warhead group. In this case, either the Leu-vinyl sulfone from EY-4-78 (resulting in 8304-vs) or the 2,4-difluorophenylalanine-epoxyketone from J-80 (resulting in 8304-ek) were synthesized. These compounds maintained potent β5 inhibition (IC_50_ = 3.5 ± 0.2 nM for 8304-vs; IC_50_ = 23.7 ± 2.1 nM for 8304-ek), similar to TDI-8304, although they also exhibited some weak inhibition of the β2 subunit in the micromolar range (**Fig. 6g-h, Table 1)**. Additionally, while the β5 subunit of human c20S was also inhibited, the selectivity for Pf20S β5 over c20S β5 was 33-fold for 8304-vs and 11-fold for 8304-ek. Importantly, these new irreversible-binding hybrid compounds demonstrated a clear reduction in the propensity to acquire resistance *in vitro* ^48^.

## Discussion

This study reports the first successful generation of reagent quantities of active, recombinant *P. falciparum* 20S proteasome by co-expressing all 14 subunits with the essential Ump1 chaperone in an insect cell system. This achievement overcomes a long-standing bottleneck in anti-malarial drug discovery research caused by the difficulty of isolating native Pf20S with high yield. The availability of a scalable and reliable source of functional Pf20S is significant because it enables high-throughput screening of compound libraries, detailed structure-function relationship studies, robust kinetic analyses of inhibitors, and advanced structure-based drug design efforts, all of which were previously challenging. This directly addresses a critical unmet need in the field, similar to how recombinant systems have advanced research on other parasitic enzymes ^33-35^.

The cryo-EM structural analysis not only validated the correctly assembled mature Pf20S but also provided new insights into the assembly process since non-catalytic immature complexes were present. This includes the visualization of Ump1 bound to half-Pf20S complexes and the distinct structural conformations of the *Plasmodium* β2 and β5 propeptides compared to their yeast or mammalian counterparts. Furthermore, the unbound β1 C-terminus in the recombinant mature form offers a molecular explanation for the incorrectly assembled proteasomes. These findings shed new light on the intricacies of proteasome biogenesis, highlighting both conserved chaperone dependencies (Ump1) and potentially unique parasite-specific assembly features or vulnerabilities. The tendency for mis-assembly, even in the presence of Ump1, suggests that *Plasmodium* proteasome assembly might have unique regulatory steps or stability determinants that differ from host or model organisms, which could be areas for future investigation and potential therapeutic targeting.

The extensive biochemical characterization confirms that the recombinant Pf20S, despite containing partially misassembled complexes, recapitulates the properties of the native enzyme. This includes the functionality of all three catalytic subunits (β1, β2, β5) with specific substrates and inhibitors, and appropriate activation by PA28α, establishing that the recombinant Pf20S is a reliable and more accessible tool than native isolates for quantitatively assessing inhibitor potency and selectivity. This allows for more standardized and reproducible assays critical for advancing Pf20S-selective inhibitors through the drug development pipeline. The identified optimal substrate panel (Ac-nPnD-amc for β1, Z-VLR-amc for β2, and Ac-FNKL-amc for β5) will be particularly useful for future screening campaigns.

The ∼3.0 Å cryo-EM structure of the potent and parasite-selective inhibitor J-80 in complex with recombinant Pf20S provides a detailed molecular understanding of its binding mode and its specificity for the β5 subunit. The structure highlights key interactions within the substrate-binding groove, particularly at the β5/β6 interface, and the formation of a morpholine ring with the catalytic threonine, revealing that the introduction of the *D*-amino acid at the P3 position leads to the change in binding pose, which may contribute to the enhanced the selectivity of Pf20S over human proteasomes. These high-resolution structural insights are invaluable for rational drug design. They provide a concrete template for optimizing J-80 or for designing new chemotypes that can achieve even greater potency and selectivity against Pf20S β5 by leveraging these specific interactions, thereby enhancing the therapeutic window for future antimalarial proteasome inhibitors.

The development and characterization of new hybrid inhibitors (8304-vs and 8304-ek), which combine the potent macrocyclic scaffold of a reversible inhibitor (TDI-8304) with irreversible warheads (vinyl sulfone or epoxyketone), represent an innovative chemical strategy. These hybrids successfully retained high potency against Pf20S β5 while demonstrating a marked reduction in the propensity to acquire *in vitro* resistance compared to the parent compound ^48^. This work highlights a promising approach to address the challenge of drug resistance, a major concern in antimalarial therapy. By converting a potent reversible inhibitor into an irreversible one while maintaining target engagement and selectivity, these hybrid compounds offer a pathway to develop more durable antimalarial proteasome inhibitors. This strategy could be more broadly applicable in the development of covalent inhibitors against other parasitic targets where resistance to reversible agents is a concern.

## Materials and methods

Bortezomib, carfilzomib, fluorogenic substrates Suc-LLVY-amc, Z-VLR-amc, Z-LLE-amc and Ac-nPnD-amc were purchased from Cayman Chemical or AdipoGen. Ac-FNKL-amc was custom-synthesized by GenScript and marizomib was purchased from MedChemExpress. The synthesis of Plasmodium-selective inhibitors is described in the relevant publications ^9,48,49^. FRSR-ek was synthesized using standard chemistry for peptide epoxyketone inhibitors. Substrates and inhibitors were dissolved in dry DMSO at a concentration of 20 or 10 mM and stored at −20°C. Human constitutive proteasome was purchased from R&D Systems, MN and stored at −80°C.

### Cloning, expression, and protein purification

Complementary DNA sequences encoding 14 proteasome subunits and the assembly chaperone Ump1 from *Plasmodium falciparum* strains Dd2 and DD7 were retrieved from PlasmoDB through sequence alignment against their human orthologs (**Fig. S1**). These sequences underwent codon optimization for *E. coli* expression systems before synthesis by Azenta Life Sciences. The optimized genes were subsequently inserted into three distinct pACEBac1 expression vectors using standard restriction enzyme cloning procedures (**File S2**). Recombinant plasmids were then introduced into DH10EmBacY competent cells (Geneva Biotech) for baculovirus generation. Recombinant protein production was carried out using the baculovirus expression system in Sf9 insect cells (ThermoFisher Scientific, catalog #11496015) following standard protocols. To evaluate the role of the Ump1 chaperone in proteasome assembly, 20S proteasome expression was conducted both with and without co-expression of this protein. Infected Sf9 cells were harvested 72 hours post-infection through centrifugation (2,000 × g, 15 minutes). Cell pellets were resuspended in lysis buffer (Buffer A: 50 mM Tris-HCl pH 7.5, 150 mM NaCl, 1 mM dithiothreitol (DTT), 1 mM EDTA) at a 4:1 volume ratio. Cell disruption was achieved through sonication, followed by clarification via high-speed centrifugation (20,000 × g, 20 minutes). The clarified cell extract was applied to tandem Streptactin XP affinity columns (QIAGEN) pre-equilibrated with Buffer A. After thorough washing, bound proteins were eluted using Buffer A containing 50 mM biotin. Proteasome-enriched fractions were pooled and concentrated via ultrafiltration (Amicon, 100 kDa) before size exclusion chromatography on a Superose 6 Increase 10/300 column (GE Healthcare). The column was equilibrated with gel filtration buffer (50 mM HEPES pH 7.5, 150 mM NaCl, 1 mM EDTA), and 0.5 ml fractions were collected for protease activity analysis. Enzymatically active fractions were combined and concentrated for further analysis.

### Proteasome activity and inhibition assays

Catalytic activity of individual proteasome subunits was assessed using subunit-selective fluorogenic peptide substrates. For human c20S proteasome, the β1, β2, and β5 active sites were monitored using Z-LLE-amc, Z-VLR-amc, and Suc-LLVY-amc substrates, respectively. The corresponding *Plasmodium falciparum* Pf20S subunits were evaluated with Ac-nPnD-amc, Z-VLR-amc, and Ac-FNKL-amc substrates. Enzymatic assays were performed in 384-well microplate format. Human c20S proteasome (1 nM) was pre-activated with 100 nM PA28α human activator, while Pf20S (1 nM) required 200 nM PA28α activator for optimal activity. The PA28α activator protein was produced recombinantly in *E. coli* using pSumo expression vector following established protocols ^50^. For c20S assays, the reaction mixture comprised of 50 mM HEPES buffer (pH 7.5) and 1 mM DTT, with fluorogenic substrates at the following concentrations: 80 μM Z-LLE-amc, 30 μM Z-VLR-amc, and 65 μM Suc-LLVY-amc. Each reaction well contained 8 μL total volume. Pf20S enzymatic reactions utilized a modified buffer system containing 50 mM HEPES (pH 7.5), 5 mM MgCl_2_, 1 mM DTT and 0.01% BSA. Substrate concentrations were optimized to 80 μM Ac-nPnD-amc, 25 μM Z-VLR-amc, and 50 μM Ac-FNKL-amc, maintaining 8 μL reaction volumes. Test compounds were precisely dispensed using an Echo650 acoustic liquid handler (Beckman Coulter). Kinetic analysis protocols were tailored to inhibitor mechanism: irreversible inhibitors underwent pre-steady-state measurements with IC_50_ determination at 60-90 minutes post-reaction initiation, while reversible inhibitors were subjected to steady-state kinetics following 30-minute pre-incubation periods. All experiments were conducted in triplicate using black 384-well microplates (Greiner Bio-One, Monroe, NC) maintained at 37°C. Fluorescence detection employed a Synergy HTX Multi-Mode Microplate Reader (BioTek, Winooski, VT) with excitation/emission wavelengths set to 360/460 nm. Data processing and statistical analysis were performed using GraphPad Prism software (version 10.4.1).

### Protein gels and active site probing

Recombinant Pf20S and Pf20S-inhibitor complex were with 2 μM Me4BodipyFL-Ahx3Leu3VS (R&D Systems #I-190). After probe addition, samples were incubated at room temperature for 16 h. For denaturing gels, samples were mixed with 4X Bolt LDS sample buffer (ThermoFisher Scientific) containing 250 μM DTT, heated at 99°C for 5 min, and loaded onto a NuPAGE 12% Bis-Tris gel (ThermoFisher Scientific). PageRuler Plus pre-stained protein ladder (ThermoFisher Scientific) was included on each gel. Gels were run with 1X MOPS SDS buffers (Invitrogen) at 120 V. For native gels, samples were mixed with 2X Novex Tris-glycine native sample buffer and loaded onto NuPAGE 3-8% Tris-glycine gels (Invitrogen) with NativeMark unstained protein standard (Thermo Fisher Scientific, 57030). Gels were run at 100 V with Novex Tris-glycine running buffer (Invitrogen). All gels were imaged on Bio-Rad ChemiDoc XRS+ at 470 nm excitation and 530 nm emission for Me4BodipyFL-Ahx3Leu3VS probe visualization and silver stained. Bands were quantified using Image Lab 6.1.

### In gel digestion and desalting

Bands for proteomic analysis were excised from a native gel and processed according to ^51^. The extracted digest was dried and resuspended in 0.1% formic acid before C18 ZipTip desalting. A C18 column was washed with methanol and spun for 45 s at 3500 × g. The column was equilibrated with 0.1% formic acid (FA) in 50% acetonitrile followed by 0.1% FA in water. The sample was loaded onto the column and centrifuged for 2 min at 2000 × g. It was then washed with 0.1% FA then eluted from C18 with 50% acetonitrile, 0.1% FA by spinning at 3500 x g for 45 s. The sample was dried in a Savant Speed Vac Plus AR and stored at −80°C.

### LC/MS analysis

LC separation was performed using a Dionex Ultimate 3000 nano HPLC system (Thermo Scientific) online coupled to the mass spectrometer. Samples were loaded onto a trap column (C18 PepMap100, 5 μm particle size, 300 μm × 5 mm, ThermoFisher Scientific) for 2 min at a flow rate of 25 μL/min. The loading buffer consisted of water with 2% acetonitrile and 0.1% trifluoroacetic acid. Peptides were separated using a linear gradient of mobile phase B from 5% to 50% over 17 min. Mobile phase A consisted of water with 0.1% FA, and mobile phase B consisted of acetonitrile with 0.1% FA. Chromatographic separation was achieved using a nano reversed-phase column (Aurora Ultimate TS, 25 cm × 75 μm ID, 1.7 μm particle size, Ion Opticks). Mass spectrometric analysis was performed on a Thermo Scientific Orbitrap Fusion mass spectrometer operating in data-dependent acquisition (DDA) mode. Eluting peptide cations were ionized by electrospray ionization in positive mode with a spray voltage of 1600 V and ion transfer tube temperature of 275°C. MS1 survey scans were acquired in the Orbitrap analyzer over the mass range of 350-1400 m/z at 120,000 resolution with the following parameters: RF lens 60%, maximum injection time 246 ms, and AGC target 1 × 10□. Precursor ions with charge states 2-7 and intensity >5000 were selected for fragmentation. Dynamic exclusion was applied for 5 s with a mass tolerance of 10 ppm. Selected precursor ions were isolated using the quadrupole with a 1.6 m/z isolation window and fragmented by higher-energy collisional dissociation (HCD) with 30% normalized collision energy. Fragment ions were detected in the ion trap with an AGC target of 1 × 10□ and maximum injection time of 35 ms. Raw MS data were processed using Thermo Proteome Discoverer software (version 3.2.0.450) with the Sequest HT search algorithm. Peptide identification was performed against the combined *Plasmodium falciparum* and *Spodoptera frugiperda* proteome database (UniProt proteome IDs: UP000001450 and UP000829999, respectively) supplemented with common contaminants (**File S1, Table S1**). Search parameters included carbamidomethylation of cysteine as a fixed modification and methionine oxidation as a variable modification. Trypsin was specified as the digestion enzyme with a maximum of 2 missed cleavages allowed. Precursor mass tolerance was set to 10 ppm and fragment mass tolerance to 0.6 Da. Protein and peptide false discovery rates (FDR) were controlled at 1% using the Percolator algorithm. Protein abundance was calculated from the summation of the peptide group abundances. Data are available via ProteomeXchange with identifier PXD067698.

### Preparation of cryo-EM grids and data acquisition

Pf20S apoenzyme (4.5 mg/mL) was prepared in 50 mM HEPES (pH 7.5) and 1 mM TCEP. For the complex, the enzyme was pre-incubated with 260 µM J-80 at room temperature for 1 h. Phosphocholine was added to a final concentration of 0.25 mM prior to cryo-EM grid preparation. Quantifoil Cu 300 mesh R2/1 grids were glow-discharged using a Pelco easiGlow system at 15 mA for 25 seconds to enhance surface hydrophilicity. 2.5 µL aliquot of the purified Pf20S or Pf20S: J-80 complex was applied to the grid, blotted for 8 seconds with a blot force of 0, using Whatman 1 filter paper at 4 °C and 100% humidity, and then plunge-frozen in liquid ethane using a Vitrobot Mark IV (ThermoFisher Scientific). Vitrified samples were imaged in a Titan Krios electron microscope (FEI/ThermoFisher) in the Cryo-EM facility of the Structural Biology Shared Resource at Sanford Burnham Prebys Medical Discovery Institute. Images were collected on a Gatan K3 detector using SerialEM ^52^ under super-resolution settings ^53^.

### Cryo-EM image processing

A total of 3,560 raw cryo-EM images were collected for the apo Pf20S sample, and 9,270 raw images were collected for Pf20S complexed with J-80. Image processing was performed separately for the apo and J-80–bound datasets using cryoSPARC (v4.7.0) ^54,55^. Motion correction of the movies was carried out using the Patch Motion Correction tool with output F-crop factor 1/2. CTF estimation was performed using the Patch CTF tool with minimum search defocus of 10,000 Å, and maximum search defocus of 30,000□Å, respectively. Initial particle picking was conducted using the Blob Picker tool, with a minimum particle diameter of 100□Å, a maximum diameter of 150□Å. The particles were extracted with a box size of 400□Å and Fourier crop to box size 100□Å and subjected to 2D classification into 50 classes. High-quality 2D classes were selected and used as templates for further particle picking. Multiple rounds of 2D classification were carried out to remove unwanted particles and enrich for well-aligned particle images. High-quality side-view 2D classes of full Pf20S were selected and re-extracted with a box size of 400 Å. Non-uniform Refinement was carried out to generate the final 3D maps.

Image processing of the half-Pf20S was performed using the Pf20S: J-80 dataset due to the larger size of this dataset compared to the apo Pf20S dataset. High-quality 2D classes of tilted half-Pf20S were selected, subjected to additional rounds of 2D classification, and re-extracted using a smaller box size of 200□Å. These particles were subsequently used to generate 3D maps using the same workflow as above.

### Model building and refinement

A high-resolution Pf20S structure (PDB ID: 7LXU) was used as an initial template to build the apo Pf20S model. The initial atomic model was fitted into the cryo-EM density map using the ‘Fit in Map’ tool in UCSF Chimera ^56^. Real-space refinement was performed residue by residue in Coot using the “real space refine zone” tool ^57,58^. The resulting model was further refined in Phenix using real-space refinement with the following parameters: maximum iterations set to 100, five macro-cycles, target bond RMSD of 0.01 Å, target angle RMSD of 1.0°, and with secondary structure restraints enabled. Model validation was conducted using the comprehensive validation tools in Phenix ^59,60^. The refinement process involved iterative rounds between Coot and Phenix to achieve satisfactory validation metrics.

To facilitate docking, J-80 was combined with the side chain of Thr_1_ from the β5 subunit as a single covalently linked structure. The J-80–Thr_1_ adduct was constructed using the Ligand Builder tool in Coot, and its corresponding geometric restraints CIF file was generated using eLBOW in Phenix with the “simple optimization” option. The J-80–Thr_1_ model was docked into the apo Pf20S structure using UCSF Chimera by aligning the Thr1 from J-80-Thr with the Thr1 of β5 subunit from the apo Pf20S. The original β5 Thr1 residue was deleted from the apo structure, and the merged J-80–Thr1 and Pf20S model was used to create the initial J-80– Pf20S model. A local refinement around the ligand binding site was performed in Coot to optimize the fit of J-80 within the cryo-EM density. Final refinement was carried out in Coot and Phenix as described above.

The apo Pf20S model served as the initial template to build the half-Pf20S structure. Fitting and refinement of this tilted half-Pf20S model were carried out in Coot and Phenix as described above.

All final cryo-EM maps and models were visualized using UCSF Chimera and ChimeraX ^61^.

## Acknowledgements

We thank and express our gratitude to Karel Harant and Jana Brezinova (IOCB, Prague) for their assistance with proteomics sample processing and mass spectrometry data acquisition.

## Data availability

The atomic models and cryo-EM maps were deposited in the Protein Data Bank and Electron Microscopy Data Bank under accession codes 9Y0L and EMD-72395 for incorrectly assembled half-Pf20S, 9Y0K and EMD-72394 for J-80 bound Pf20S, 9Y1O and EMD-72400 for apo Pf20S. Source data for the cryo-EM map density analysis are provided in the Source data file with this paper. The mass spectrometry proteomics data have been deposited to the ProteomeXchange Consortium via the PRIDE^62^ partner repository with the dataset identifier PXD067698 and 10.6019/PXD067698

