## Supplementary material for "A recombinant expression system for the *Plasmodium falciparum* proteasome enables structural analysis of its assembly and the design of selective inhibitors": rPf20S_Supplemental_Fajtova.docx

**Supplementary Figures**

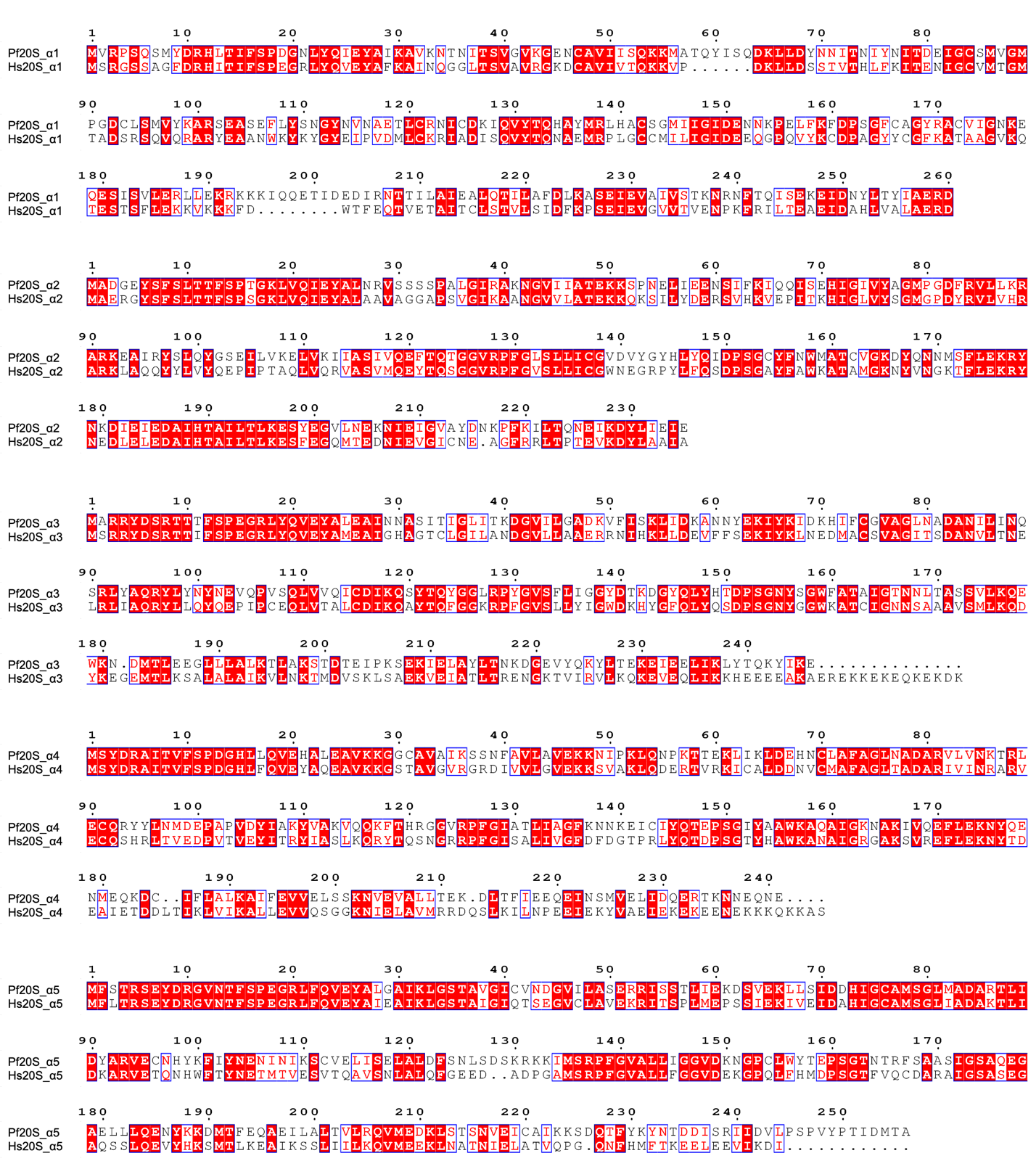

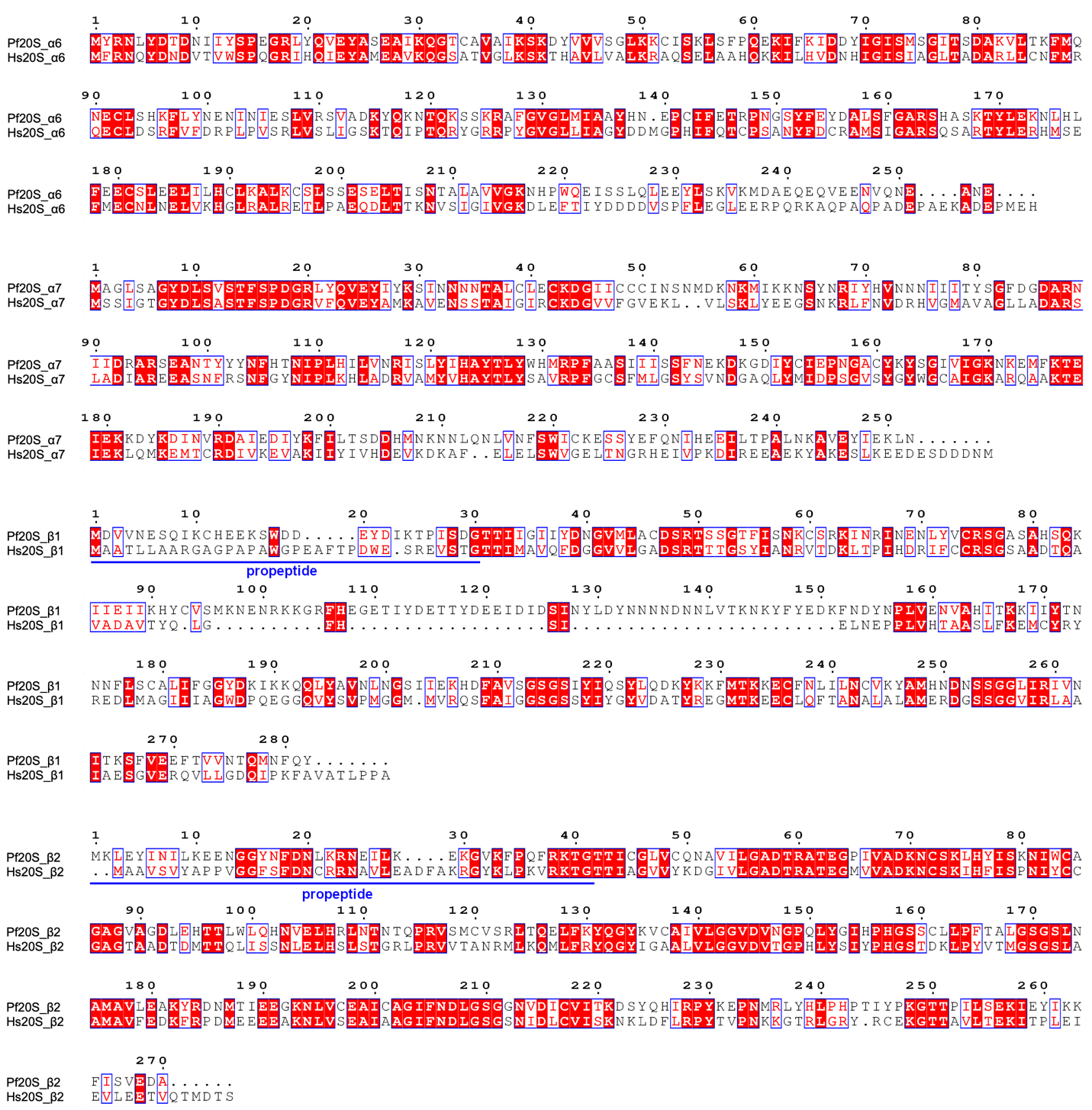

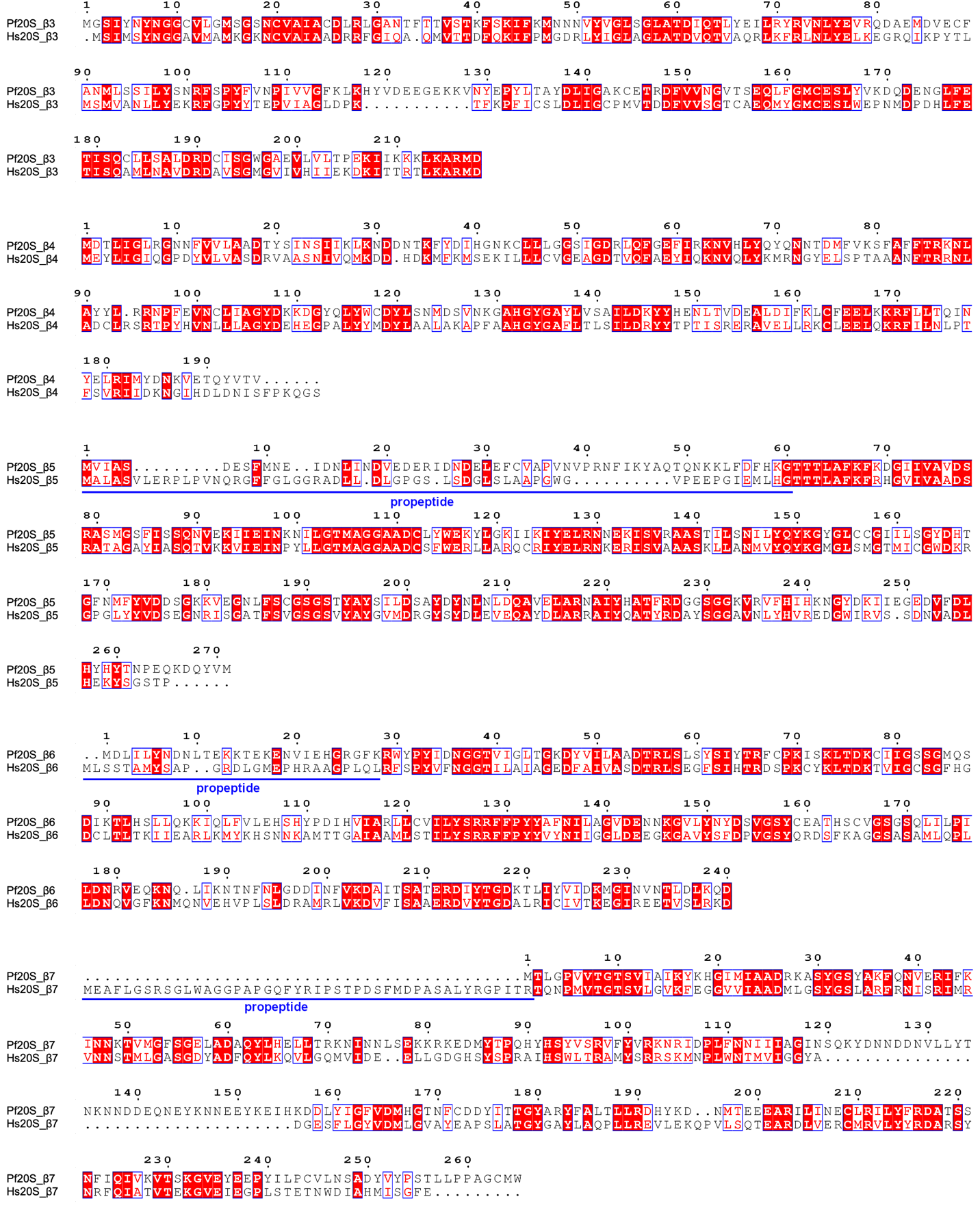

**Supplementary Figure 1** Alignment of seven α and seven β subunits of *Plasmodium falciparum* proteasome (Pf20S) with the homologous subunits in the *Homo sapiens* 20S proteasome, commonly known as the constitutive proteasome (c20S). The aligned sequences were visualized using ESP3 software according to Robert, X. and Gouet, P. (2014).

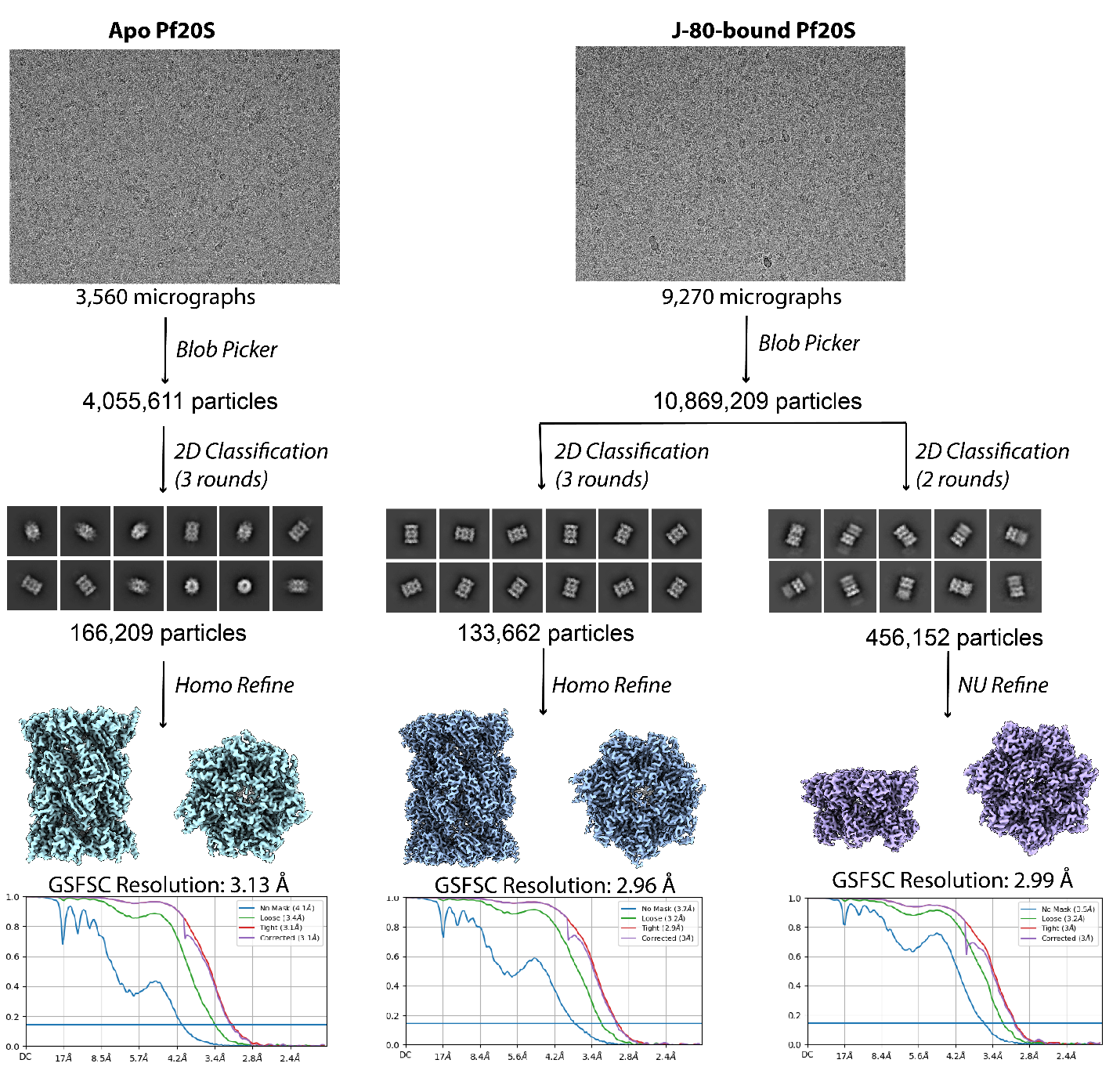

**Supplementary Figure 2.** Cryo-EM workflow of data processing. This image processing workflow was employed to reconstruct the Pf20S structures in cryo-EM. Homo Refine and NI Refine and GSFSC correspond to Homogeneous Refinement, Non-uniform Refinement, and Gold-Standard Fourier Shell Correlation, respectively.

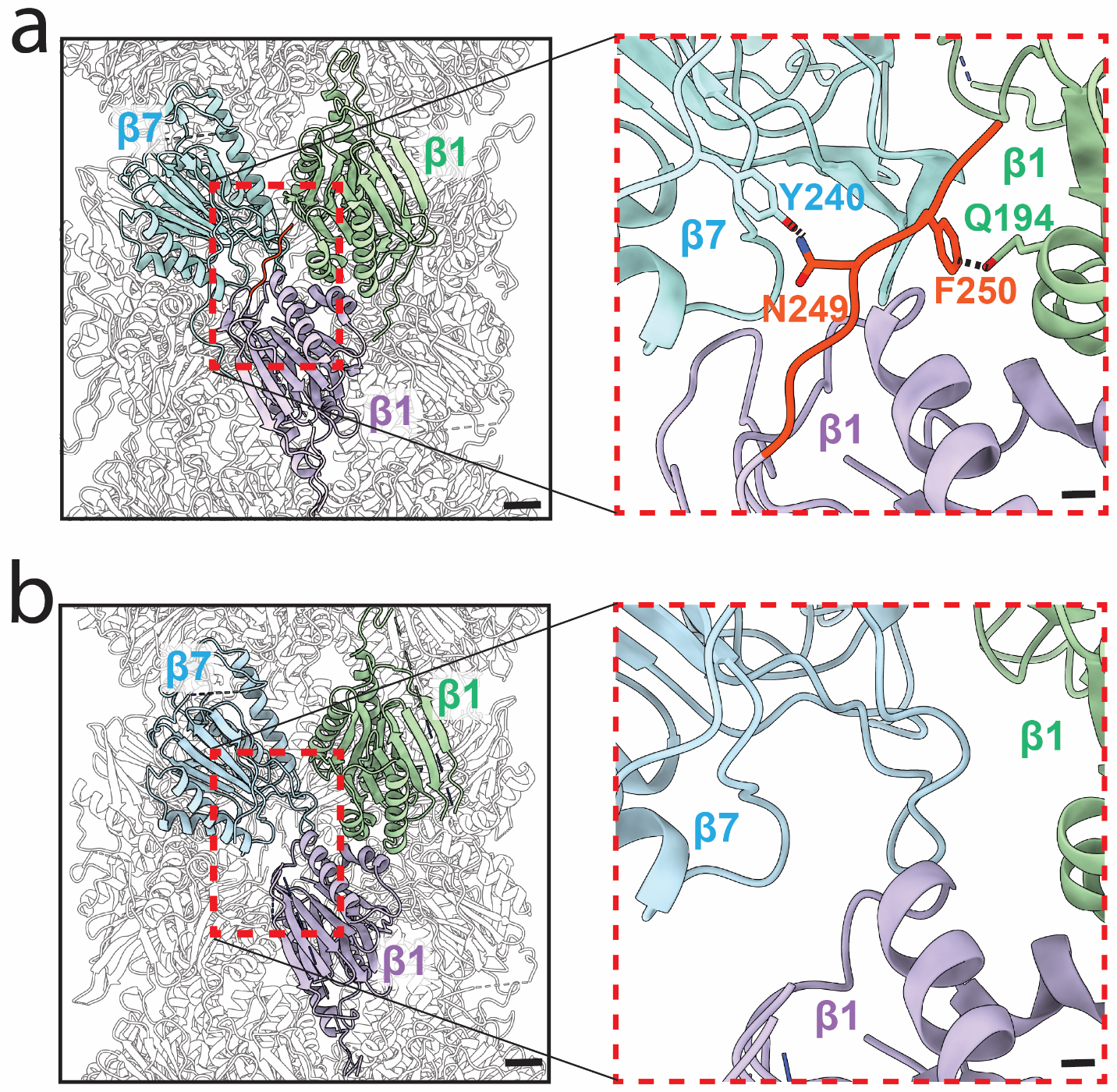

**Supplementary Figure 3. C-terminal of β1 is not observed in the recombinant Pf20S.** **a**. In the endogenously purified Pf20S structure (PDB: 7LXU), the β1 C-terminus (red box) extends across the dimerization interface of the half-proteasome and is positioned between β7 and β1 of the opposing half-Pf20S. **b**. The β1 C-terminus is not observed in the recombinant Pf20S. Scale bars: 20 Å for left panels, 5 Å for right panels.

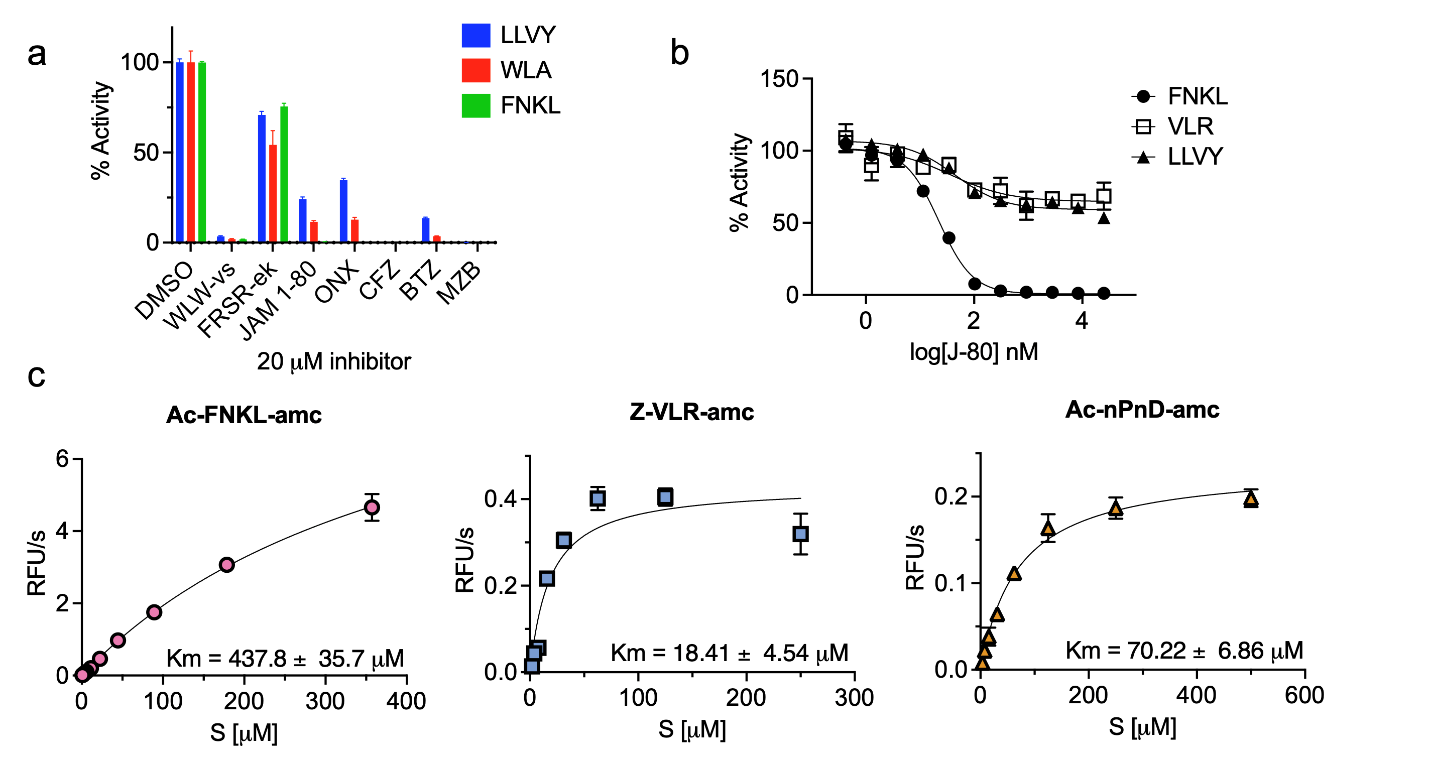

**Supplementary Figure 4. Discovery of new Pf20S β5 reporter substrate and Km studies** a. Pf20S preincubated with a set of subunit selective inhibitors. The enzyme activity was monitored using Suc-LLVY-amc, Ac-WLA-amc and Ac-FNKL-amc substrates. b. Concentration response using J-80 and Pf20S c. Michaelis-Menten saturation curve for the three subunit-specific substrates.

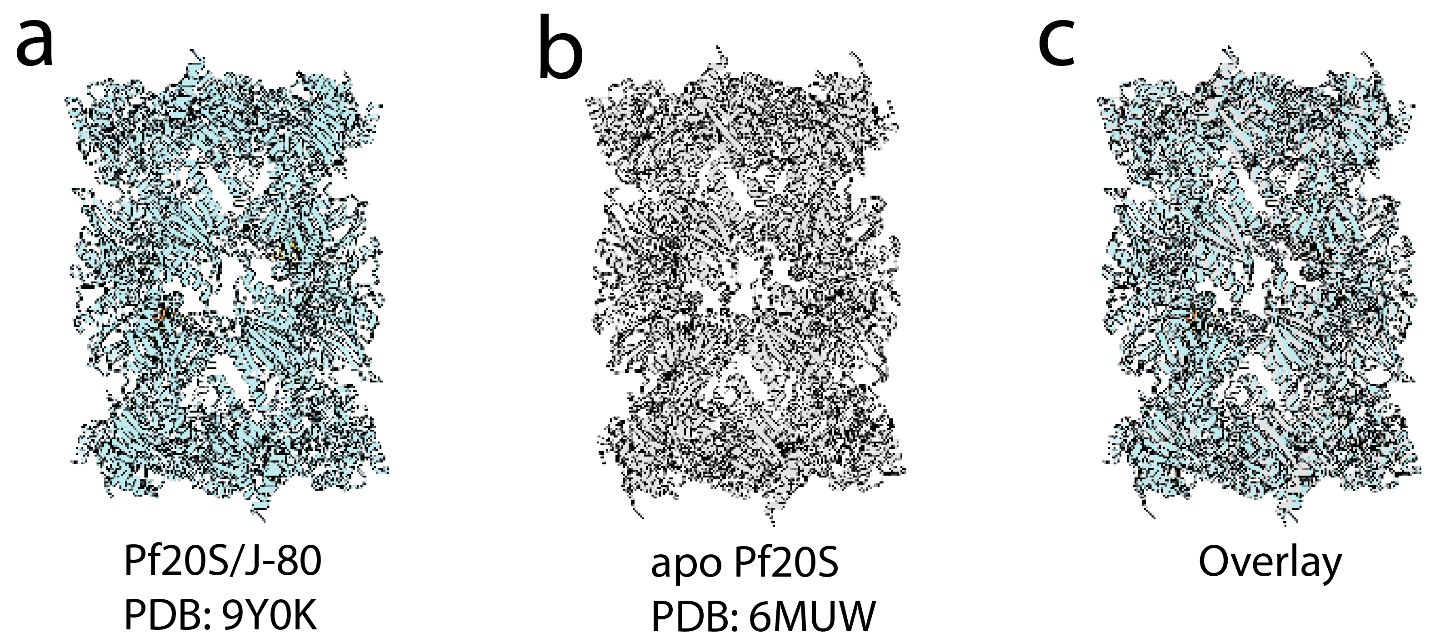

**Supplementary Figure 5**. Structural comparison of the apo form versus the Pf20S/J-80 complex. The root-mean-square deviation (RMSD) calculated over 230 pruned atom pairs is 0.78 Å, indicating minimal conformational change between the two states, highlighting strong structural consistency.

**Supplementary Tables**

**Supplementary Table 1** Uniport access numbers

| **Subunit** | **Hs Uniprot ID** | **Sf Uniprot ID** | **Pf Uniprot ID** |
| --- | --- | --- | --- |
| **α1** | [P60900](https://www.uniprot.org/uniprotkb/P60900) | A0A2H1W7W9/A0A9R0EML5 | Q8IAR3 |
| **α2** | [P25787](https://www.uniprot.org/uniprotkb/P25787) | A0A2H1VM92 | C6KST3 |
| **α3** | [P25789](https://www.uniprot.org/uniprotkb/P25789) | A0A2H1WB52/A0A9R0DNT9 | Q8IDG3 |
| **α4** | [O14818](https://www.uniprot.org/uniprotkb/O14818) | A0A2H1WH41 | Q8IDG2 |
| **α5** | [P28066](https://www.uniprot.org/uniprotkb/P28066) | A0A2H1VS21 | [Q8IBI3](https://www.uniprot.org/uniprotkb/Q8IBI3/entry) |
| **α6** | [P25786](https://www.uniprot.org/uniprotkb/P25786) | A0A1S6Q5K8/A0A9R0D7L3 | [Q8IK90](https://www.uniprot.org/uniprotkb/Q8IK90/entry) |
| **α7** | [P25788](https://www.uniprot.org/uniprotkb/P25788) | A0A2H1X1L9 | [O77396](https://www.uniprot.org/uniprotkb/O77396/entry) |
| **β1** | [P28072](https://www.uniprot.org/uniprotkb/P28072) | A0A2H1VFD5 | Q8I0U7 |
| **β2** | [Q99436](https://www.uniprot.org/uniprotkb/Q99436) | A0A2H1WXA3 | [Q8I6T3](https://www.uniprot.org/uniprotkb/Q8I6T3/entry) |
| **β3** | [P49720](https://www.uniprot.org/uniprotkb/P49720) | A0A2H1VUK5 | Q8I261 |
| **β4** | [P49721](https://www.uniprot.org/uniprotkb/P49721) | A0A2H1VJ26 | Q8IKC9 |
| **β5** | [P28074](https://www.uniprot.org/uniprotkb/P28074) | A0A2H1W5W8 | Q8IJT1 |
| **β6** | [P20618](https://www.uniprot.org/uniprotkb/P20618) | A0A2H1W3L5 | A0A5K1K7U1 |
| **β7** | [P28070](https://www.uniprot.org/uniprotkb/P28070) | A0A2H1W6J7 | Q7K6A9 |
| **Ump1** | [Q9Y244](https://www.uniprot.org/uniprotkb/Q9Y244) | A0A2H1X3E3 | A0A5K1K941 |

**Supplementary Files**

**Supplementary File 1**. Proteomic data showing of upper and lower bands excised from gel and searched against the *P. falciparum* and *S. frugiperda* proteomes.

**Supplementary File 2**. Sequence of pACEBac1 expression vectors
